# Parameter exploration improves the accuracy of long-read genome assembly

**DOI:** 10.1101/2021.05.28.446135

**Authors:** Anurag Priyam, Alicja Witwicka, Anindita Brahma, Eckart Stolle, Yannick Wurm

## Abstract

Long-molecule sequencing is now routinely applied to generate high-quality reference genome assemblies. However, datasets differ in repeat composition, heterozygosity, read lengths and error profiles. The assembly parameters that provide the best results could thus differ across datasets. By integrating four complementary and biologically meaningful metrics, we show that simple fine-tuning of assembly parameters can substantially improve the quality of long-read genome assemblies. In particular, modifying estimates of sequencing error rates improves some metrics more than two-fold. We provide a flexible software, CompareGenomeQualities, that automates comparisons of assembly qualities for researchers wanting a straightforward mechanism for choosing among multiple assemblies.

## Background

High-quality genome assemblies are essential for modern biological research. Genome assemblies serve as the reference for integrative study of organismal biology [1,2] and for phylogenomic comparisons [3,4]. Unfortunately, eukaryotic genome assemblies typically contain major errors. This is because eukaryotic genomes include large amounts of repetitive sequences that are difficult to resolve due to the limitations of sequencing processes and assembly algorithms [5]. The inability to resolve repetitive sequences leads to assembly fragmentation [6], to collapsing of multiple occurrences of repetitive sequence into fewer assembled sequences [7], and to misassembly of repetitive regions [8]. Such shortcomings of genome assemblies reduce the sensitivity and specificity of downstream analyses. For example, assembly fragmentation can lead to underestimation of syntenic relationships [9], and errors in gene prediction [7,10,11]. Furthermore, when sequence reads from different copies of a repetitive element map to a collapsed representation of the repeat, small differences between the repeat copies can be incorrectly identified as polymorphisms [12].

Long-molecule sequencing can dramatically improve genome assemblies [13]. In particular, long reads can span tandem arrays of repetitive elements or interspersed repeats and thus help to resolve their sequences and structures [14]. Furthermore, long-molecule sequencing technologies are more robust to variation in GC composition than short-read technologies [15]. However, good data alone cannot guarantee a good assembly. The ability of assembly software to reconstruct the correct genome sequence varies across species, sequencing technologies, and algorithmic parameters [16–20]. This suggests that *de novo* genome assembly projects are likely to benefit from testing different assembly software and algorithmic parameters for their datasets. This requires overcoming two associated challenges: which algorithmic parameters to optimize, and how to compare assemblies in order to identify the best one.

Assemblers can have dozens of parameters, making an exhaustive search of the parameter space of most assemblers impractical. However, the central principle of genome assembly software is to determine overlaps between pairs of reads and stitch together reads that overlap the best [21]. Changing the parameters that impact the read overlapping process should thus have substantial impact on assembly quality. Indeed, for the popular Canu and FALCON assemblers, modifying minimum read length and minimum overlap length parameters can improve assembly quality [18]. Another parameter that should similarly affect assembly quality is the estimate of sequencing error used by the assembly software. If the true sequencing error rate is higher than the estimate used by the software, then true overlaps between reads would be missed. This would fragment the assembly. Alternatively, if the true sequencing error rate is lower than the estimate used by the software, the number of false overlaps would increase. This can lead to assembly fragmentation, collapse, or mis-assembly of repetitive regions.

An assembly is better if it is more contiguous, accurate, and complete. The N50 length, which indicates that 50% of the assembled genome is in pieces longer than N50, provides a useful view of contiguity even if not biologically meaningful. In contrast, testing for the presence and completeness of protein-coding genes from related organisms [22] or concordance with transcriptomic data [10,23] can indicate assembly accuracy and completeness, but only in genic regions. Genome-wide measures of completeness or accuracy are less apparent. Many current projects lack datasets that would be ideal for such comparisons, including sequences from independent fosmid or BAC libraries, high-resolution genetic, optical, or chromatin interaction maps, or a high-quality reference assembly. Independently derived pairs of short Illumina DNA sequences exist for most long-molecule genome projects and these short reads can be used to detect structural errors in an assembly [24] or provide a base-by-base view of consensus accuracy [25]. Appropriately combining information from different quality metrics could provide a holistic view of genome assembly quality. However, the efforts required to identify the most meaningful metrics, collecting these metrics for multiple assemblies, and deriving summary statements of assembly accuracy and completeness requires considerable efforts.

To test the impact of varying the estimates of sequencing error on assembly quality and to establish a simple approach for selecting the best assembly, we obtained Pacbio reads for the red fire ant, *Solenopsis invicta* and generated 36 assemblies using Canu [26]. This species is a model for the study of social behavior, and a globally invasive pest [27]. The draft genome assembly for this species [28] has been cited more than 350 times despite its high fragmentation (69,511 sequences) and capturing only 79% of the genome [29]. Importantly, the fragmentation and the missing sequences affect genomic regions involved in environmental perception [30,31], and complex behavioral and developmental traits [32–36]. To compare the generated assemblies, we used four complementary metrics that characterize assembly contiguity, completeness, and accuracy. We show that varying error thresholds for finding overlaps between reads significantly improves contiguity, completeness, and accuracy of Canu assemblies. We present a tool that enables other researchers to easily compare and rank assemblies.

## Results

### Pacbio dataset and assembly parameters

We obtained 2.9 million Pacbio reads, totaling 20.2 billion bases (45x genome coverage) from a diploid sample of *S. invicta* (N50 read length of 8,876 bp; Figure S1, Additional file 1). We first assembled this dataset using default parameters of Canu. We then generated 35 additional assemblies to test the effects of three parameters (full details in Table S1). We varied the raw overlap error rate threshold, using values corresponding to sequencing error rates of 12.5%, 13.75%, 15% (default), 16.25%, and 17.5%. We varied the stringency of trimming raw reads, requiring a minimum of 4 overlaps (default), a more relaxed setting of 2 overlaps, and disabling trimming of raw reads altogether. We included this parameter in our tests because the default read trimming setting resulted in eliminating a relatively high amount, 28%, of our raw data. Finally, we varied the overlap error rate threshold for “corrected reads” that are generated by Canu at the end of the first step of the assembly pipeline. We tested values corresponding to sequencing error rates between 1.15% and 5.87% (default: 2.25%).

Long-read genome assemblies can contain considerable residual sequencing errors and unresolved haplotypes, *i*.*e*., genomic segments represented more than once in the assembly, typically due to high divergence or structural differences between haplotypes present in the original sample. To minimize the impacts of such issues on comparisons of assembly qualities, we performed one round of assembly “polishing” [37] and unresolved haplotype removal [38] prior to calculating assembly quality metrics.

### Measures of assembly contiguity, accuracy and completeness

To compare the 36 genome assemblies, we obtained four metrics of assembly quality. We first calculated the NG50 metric, which is the N50 metric normalized by estimated genome size. Second, we determined the BUSCO score, which is the number of expected single-copy genes (n=4,415) present and intact in the assembly [22]. Third, we obtained and mapped short-read Illumina sequences from a PCR-free sequencing library to each assembly. This mapping enabled us to measure the resolved length of each assembly, which we defined as the cumulative length of the regions that have between 5x coverage and twice the median coverage (Figure S2A, Additional file 1). Instead of total assembly length, which can be affected by assembly artifacts, the resolved length metric shows how much of the genome is potentially usable for analysis through standard approaches. The lowest-coverage regions can be symptomatic of sequencing or assembly issues. Similarly, regions with particularly high coverage typically contain collapsed repeats and cause false-positives in SNP datasets. Finally, we measured the percentage of solid read pairs, which we define as the percentage of all read pairs that mapped in their entirety (*i*.*e*., without clipping) and within the expected distance and orientation of their mate (*i*.*e*., concordantly) to resolved regions of the genome. This metric summarizes assembly accuracy because assembly errors such as mis-joins, inversions, collapses and consensus errors often cause clipped and non-concordant read mapping [39]. This metric also summarizes assembly completeness, as all reads are expected to map to a complete assembly of the organism’s diploid genome. Furthermore, unlike likelihood-based metrics of assembly quality [40], the absolute value of solid read pairs is meaningful in its own right as it should approach 100% for perfect sequences and a perfect assembly.

### Four complementary metrics reveal extensive variation in assembly quality

We found a 2.3-fold difference in the NG50 metric of contiguity between assemblies (237,734 bp to 543,457 bp). We similarly found 1.4-fold variation in the number of missing or incomplete single-copy genes (141 to 202). Furthermore, resolved assembly lengths vary up to 12.6 Mb, *i*.*e*., by up to ∼2.8% of genome size. Finally, there was a 2.6% range in the proportion of Illumina read pairs that map concordantly to resolved regions of the assemblies. These four measurements of assembly quality have positive but weak correlations (average 0.66; Spearman’s rank correlation coefficient), highlighting their complementarity and the importance of considering multiple measures of genome quality (Figure S3, Additional file 1).

To select the best assembly, we summed the ranks of the assemblies in each metric, weighted by the complement of the average correlation of the metric with other metrics (Fig. 1). Twenty-three assemblies (64%) had higher overall quality than obtained through default parameters. In particular, the best ranked assembly had 17.2% higher NG50 (518,074 vs 441,945 bp), 11.3% less missing or incomplete expected single-copy genes (141 vs 159), 1.8 Mb higher resolved length and 0.33% more solidly mapping Illumina reads (57.81% vs 57.62%) than the default assembly. This best ranked assembly was based on an overlap error threshold corresponding to a sequencing error rate of 13.75% for raw reads, 3.45% for corrected reads, and no trimming of raw reads.

**Fig. 1:**
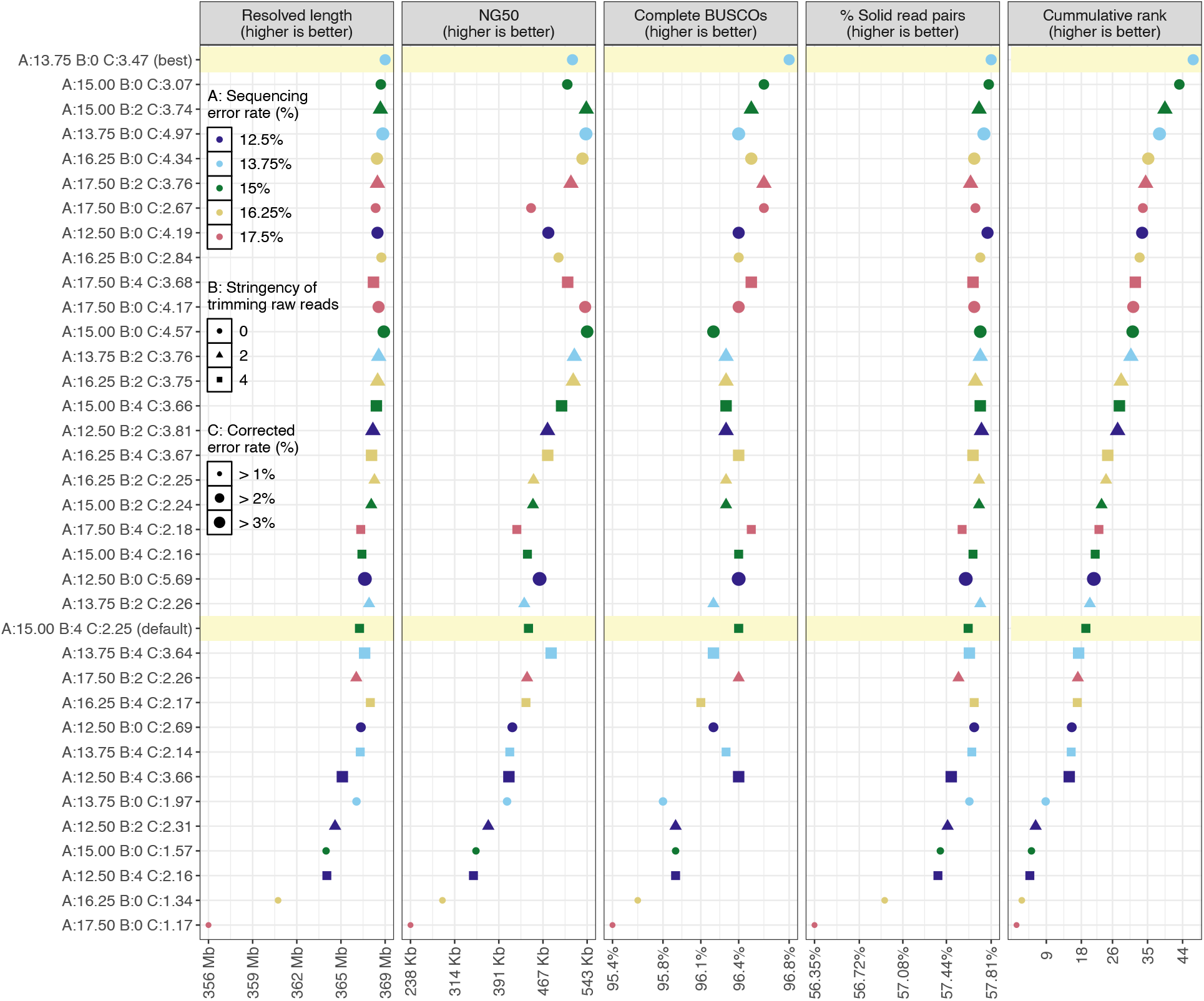
Assembly qualities and overall rank. Thirty-six polished genome assemblies ordered on the y-axis from best (top) to worst (bottom) based on the weighted sum of their ranks (rightmost panel) in each of the four metrics (other panels). The x-axis shows the range of values of each metric, or of the weighted rank in case of the rightmost panel. The best assembly and the one generated using default parameters are highlighted in yellow. Twenty-three assemblies scored higher than the ‘default’ assembly. This visualization is derived from the output of our CompareGenomesQualities tool.

In this experiment, the estimated error rate for corrected reads had the most significant impact on the overall assembly quality (generalized linear model; p < 10^−5^), followed by the estimated error rate for raw reads (p < 0.05). There was no general trend for the impact of raw read trimming on assembly quality (p = 0.5).

### Processing and chromosome-level scaffolding of best assembly for use as the reference assembly

To make the best assembly suitable for use as the reference assembly for the red fire ant, we corrected residual sequencing errors [41] and replaced rare alleles in the assembly [42] by mapping short read population-sequencing datasets (270x genome coverage) [29,36] to the assembly and substituting the most common variant at each locus [43]. Additionally, we removed contigs that appeared to be from bacteria, fungi or plants (Table S2). Finally, we ordered and oriented the contigs into chromosome-level scaffolds using genetic maps, complemented by optical maps, and paired RNA sequencing reads. The resulting assembly captures 347 Mb (77%) of the fire ant genome in 16 chromosomes, and another 38 Mb (8%) in 916 unplaced contigs (Figure S2B, Additional file 1). At time of writing, this is the most complete genome assembly of the red fire ant *Solenopsis invicta* (Table S3). A comparison with the draft genome of the species [28] shows high collinearity and the inclusion of many more sequences in our assembly (Figure S4, Additional file 1).

## Discussion

We show that small changes in the estimates of sequencing error rates used by the genome assembly software Canu produced more contiguous, accurate and complete assembly of the red fire ant genome than when using default parameters. The best assembly was obtained by lowering the estimated sequencing error rate for raw reads but increasing the estimated error rate for corrected reads. The first change suggests that default parameters may lead to false-positive detection of overlaps – probably among copies of repetitive sequences – and erroneous reciprocal correction of the repeat copies. The second change suggests that read correction does not always live up to the standards expected by subsequent steps of the assembly software. The general message that changing parameters affects outcomes should hold for other datasets. However, the impacts of particular parameter levels will depend on dataset specific features including repeat composition of the genome and the lengths and the error profiles of sequenced reads. For example, to obtain the highest quality assembly from a different Pacbio dataset from the same species, we had to increase the overlap error rate thresholds for raw reads by 2% and decrease error rate estimates for corrected reads by 0.5% (data not shown).

Our work also shows the importance of considering multiple metrics that can reveal independent aspects of assembly quality. To fill a gap in existing metrics, we also estimated genome-wide assembly completeness and accuracy using a new metric, the percentage of solidly mapping Illumina read pairs. The idea behind the metric is two-fold. First, a completely resolved assembly should have a near homogenous coverage. This is because if all copies of a repeat are present in the assembly, the read mapper will distribute the reads evenly across the different copies even if it cannot precisely determine which copy the read originated from. Additionally, if the assembly accurately represents the genome, all reads should map in their entirety to a contiguous stretch of the assembly (*i*.*e*., without clipping or splitting of reads, and concordantly with respect to their mate). This only holds if Illumina read pairs are derived from the same individual that was sequenced for genome assembly. Furthermore, contaminants, mapping errors and the haploid nature of genome assemblies means that not all reads will map perfectly to a perfect assembly. However, a higher value of the metric should indicate a more complete and accurate assembly. Because of this simplicity of interpretation and simultaneous quantification of both completeness and accuracy, we expect that our metric can become a standard for reporting the quality of published genome assemblies alongside N50 and BUSCO.

Lastly, rather than linearly combine the results of different assembly metrics, which can overemphasize correlated characteristics, we weight the metrics by their relative independence. This provides a robust framework for comprehensive comparison of assembly qualities. To simplify the application of our genome comparison approach we have created a tool, CompareGenomeQualities, that will derive the four complementary metrics we presented and rank the assemblies based on weighted sum of ranks, producing summary tables and figures analogous to Fig. 1. The tool is agnostic to the assembly approach: as inputs it requires a set of genome assemblies in FASTA format, an estimated genome size, indication of which taxonomic phylum to use for BUSCO score calculations, and paired Illumina sequences (Supplementary text, Additional file 1). Furthermore, the tool is modular, thus additional metrics of assembly quality, such as those obtained from QUAST [16] or Merqury [44], can be included for ranking and visualization using simple tabular files.

## Conclusions

We show that tweaking algorithmic parameters used by genome assembly software can significantly improve assembly qualities. In particular, we find that the estimates of sequencing errors used by assembly software are relevant parameters to optimize. Furthermore, given the challenges of considering biologically relevant metrics of genome quality to compare genome assemblies, we present a tool, CompareGenomeQualities, that automates this process. The tool combines complementary metrics of contiguity, completeness, and accuracy. Contiguity is measured by normalizing the classic N50 metric by genome size. Completeness and accuracy are measured in genic regions by testing for the presence of expected single-copy genes (BUSCO score) [22], and of the whole genome using two metrics derived from mapping characteristics of Illumina reads. We expect that CompareGenomeQualities will be helpful to the many researchers now sequencing eukaryotic genomes.

## Methods

### Sample collection and sequencing

We collected male pupae of the fire ant *Solenopsis invicta* from one single-queen colony from Campo Grande, Brazil (GPS coordinate: 20°38’46.85”S 50°38’36.58”W, permit number: 14BR015531/DF). Since the pupae are from a single-queen colony, they are full brothers. Males of this species are haploid, while the females are diploid. Samples were flash-frozen and preserved at -80° centigrade until further processing. Species and colony organization (*i*.*e*., single- or multiple-queen) were confirmed respectively using partial sequencing of the mitochondrial cytochrome c oxidase I gene and an RFLP marker assay [29].

### Pacbio sequencing of a pool of 21 haploid brothers for assembly

We extracted DNA from twenty-one pupae using a CTAB-phenol-chloroform protocol [45]. From this DNA, the Centre for Genomics Research in Liverpool prepared an SMRT library with a size selection of 10 kb and sequenced the library using 5 SMRT cells on a Pacbio Sequel device (V2 chemistry).

### Assembly parameters and workflow

We generated a total of 36 assemblies from the Pacbio sequences, using Canu (version 1.6) [26]. We generated one assembly using default parameters to serve as a reference point for all comparisons. We generated the remaining 35 assemblies to test the effects of three parameters: error rate threshold for detecting overlaps between raw reads (rawErrorRate), minimum number of overlaps required to not trim or split raw reads (corMinCoverage), and error rate threshold for detecting overlaps between corrected reads (correctedErrorRate). For rawErrorRate, we tested the values 0.25, 0.275, 0.30 (default), 0.325, and 0.35 corresponding to sequencing error rates of 12.5%, 13.75%, 15% (default), 16.25%, and 17.5%. For corMinCoverage, we tested the values 4 (default), 2, and 0. Zero disables trimming and splitting of raw reads, whereas two represents a more lenient trimming and splitting stringency compared to the default. For correctedErrorRate, we tested values specific to each combination of rawErrorRate and corMinCoverage. That is, we used the -correct option of Canu to generate corrected reads for the fifteen combinations of rawErrorRate and corMinCoverage. We then estimated the error rate of corrected reads by mapping them to the GCF_000188075.1 assembly [28] using minimap2 (version 2.5-r574; -a -L -x map-pb) [46] and calculating the total edit distance between the reads and the reference divided by the total number of mapped bases (Figure S5, Additional file 1). We only considered coding regions of highly conserved, single-copy genes for the calculation (n=988), because reads mapping to such regions are extremely unlikely to be mismapped. The gene structures were downloaded from Ensembl BioMart, those matching the criteria: orthologous to the nematode *C. elegans* and without a paralog. We derived the mismatch rate by obtaining a pileup of the primary alignments in the coding regions of the genes using samtools (version 1.4.1) [47]. The fifth column of the pileup format provided the number of mismatches, and the fourth column provided the number of mapped bases. At first, we set correctedErrorRate to twice the estimated error rate and generated one assembly for each of the 15 combinations of rawErrorRate and corMinCoverage. However, ten out of the fifteen assemblies were highly fragmented (N50 < 100 kb), suggesting more noise in corrected reads than estimated. Indeed, for the set of corrected reads obtained using default parameters, our estimate of the error threshold deviated from the default value by almost 3%. We thus assembled each set of corrected reads twice more by increasing the calculated error threshold by 3% and by 6% and generated 30 more assemblies. Overall, we tested error rate of corrected reads between 1.15% and 5.87%. We excluded the ten assemblies that had N50 lower than 100 kb from comparisons.

For all except the default assembly, we changed two other parameters from their default values. By default, Canu’s read correction step only corrects the longest input reads that would represent 40x genome coverage. However, as trimming of raw reads alone (corMinCoverage) can discard up to 28% of data, we were apprehensive of losing more and disabled further subsetting of input reads by setting corOutCoverage to 100 [48]. Additionally, we changed the corMhapSensitivity parameter from “normal” to “high” to increase the sensitivity of overlap detection between raw reads [49].

We polished all assemblies and removed “unresolved haplotigs” before comparison as they can impact BUSCO and read mapping metrics (Table S4). For polishing, we used raw Pacbio data in BAM format with the SMRTLink software suite (version 5.1.0.26412), which takes into account quality signals inherent to SMRT sequencing [37]. To remove unresolved haplotigs, we used Pacbio reads in FASTA format with the purge_haplotigs pipeline (version 0b9afdfd) [38], which works on the principle that redundantly assembled loci will have high sequence similarity and half the mean genome coverage. Minimap2 (version 2.5-r574; -a -L -x map-pb) [46] was used to map Pacbio reads to the assemblies; we discarded reads shorter than 1000 bp before mapping. Figure S6 (Additional file 1) shows coverage histograms of the best assembly before and after running purge_haplotigs. The best assembly was further polished using Illumina reads later (see below, “Removal of residual sequencing errors and rare alleles from the best assembly”).

### Assembly quality metrics and ranking

For each assembly, we obtained measures of contiguity, accuracy and completeness. First, we used QUAST (version 4.6.1) [50] to get the NG50 metric of contiguity assuming the genome size to be 450 Mb [29]. Second, we used BUSCO (version 3.0) [22] to determine how many of the genes expected to be present in a single copy in Hymenopteran species (n=4,415) are indeed present and intact in each assembly. This BUSCO score provides a measure of assembly accuracy and completeness in genic regions. For a genome-wide measure of accuracy and completeness, we downloaded Illumina read-pairs derived from a brother of the individuals used for Pacbio sequencing and from another male of a nearby colony: SRA runs SRX4907869 and SRX4907871, respectively [29]. We cleaned the Illumina reads (Supplementary text, Additional file 1) and mapped them to the assemblies using default parameters of bwa-mem (version 0.7.17) [51]. Next, for each assembly, we used mosdepth (version 0.2.6) [52] to obtain read depth at each base (or, 1 bp windows) of the assembly in a BED file. We then filtered the windows with depth lower than 5x (assembler chaff) or higher than twice the median coverage (collapsed regions) using custom scripts. The number of bases retained after filtering is the resolved length of the assembly, a measure of assembly completeness. Next, we used bedtools (version 2.28.0) [53] to obtain the subset of Illumina read mappings that overlapped with resolved regions of the genome. Finally, using a custom script, we counted the number of Illumina read-pairs that overlapped with resolved regions of the genome and mapped such that neither read of the pair was clipped and both the reads mapped concordantly with respect to each other. The read-pairs that fulfill the above criteria are considered to be solidly mapped and provide a measure of assembly accuracy and completeness.

To consolidate the four assembly quality metrics into an overall rank, we first ranked the assemblies by each metric. We then calculated Spearman’s rank correlation coefficient between pairs of metrics and, from this, each metric’s average correlation with all other metrics. Finally, we summed the ranks of the assemblies, weighted by one minus the average correlation of the metric with other metrics (*i*.*e*., the complement of the average correlation).

### Determining the significance of assembly parameters

We modelled the overall assembly rank as a function of the three assembly parameters (Figure S7, Additional file 1). Interaction terms were removed from the model in a stepwise procedure based on their level of significance. To ensure the data fit the assumptions of the linear model, we inspected homoscedasticity, multicollinearity, the relationship between residuals and predicted values, and recognized them as satisfactory across the model.

### Removal of residual sequencing errors and rare alleles from the best assembly

To remove residual sequencing errors and rare alleles from the best assembly, we used eighteen Illumina whole-genome sequence datasets along with the Pacbio reads used for assembly. The Illumina datasets included all thirteen “bigB” labelled SRA runs from BioProject PRJNA542606 [36], and all five “bigB” labelled SRA runs from BioProject PRJNA396161 [29]. We cleaned the Illumina reads (Supplementary text, Additional file 1) and mapped them to the assembly using default parameters of bwa-mem (version 0.7.17) [51]. We mapped the raw Pacbio reads to the assembly using minimap2 (version 2.17; -a -L -x map-pb) [46]; we discarded reads shorter than 1000 bp before mapping. Finally, we used pilon (version 1.23) [43] on the assembly and the resulting alignments to generate a polished assembly. Here, Pacbio sequences are used to disambiguate Illumina read mappings in repetitive regions of the genome [54].

### Identification of foreign DNA in the best assembly

To identify foreign DNA in the best assembly, we used Kraken2 (version 2.0.8) [55] to compare the contigs to NCBI’s non-redundant databases of nucleotide sequences (downloaded on April 22, 2020) and 231 new, insect viral sequences from the literature [56].

### Ordering and orienting contigs

To assign the polished and filtered contigs to one of the sixteen fire-ant chromosomes, we generated genetic maps from RAD sequencing (RADseq) of seven fire ant families [32]. We further derived contig connectivity information from Bionano optical maps [29] and RNA sequencing (RNA-seq) of various tissue types and developmental stages: all SRA runs from BioProjects PRJNA542606 [36], PRJNA422376 [57], PRJNA266847, and PRJNA393960. We provided these as input to ALLMAPS (version 0.8.12) [58], assigning them equal weight to reduce the propagation of biases of any one dataset.

To create genetic maps, we first demultiplexed the RADseq reads using a custom script and cleaned the demultiplexed reads using default parameters of stacks2 (version 2.5) [59]. Second, for each family, we mapped the cleaned RADseq reads to the assembly using default parameters of bwa-mem (version 0.7.17) [51] and genotyped the individuals using stacks2 (-X “populations: -e ecoRI --vcf”). The VCF output of stacks contained only bi-allelic sites. Next, for each family, we plotted the number of called sites for each individual on the x-axis and the corresponding number of homozygous sites on the y-axis (Figure S8, Additional file 1). Because the individuals are haploid, we expect an almost 1:1 correlation between the number of called sites and the number of homozygous sites. We performed a linear regression in R (y ∼ x + 0) and eliminated individuals that were two standard deviations away from the regression line. We additionally removed individuals that jumped out on the plot as having too few called sites.

Next, we filtered variant sites based on the number of missing observations (because the individuals are haploid males, we treated heterozygous calls as missing observation), mean site depth, mean genotype quality, and minor allele frequency. The respective thresholds were chosen by inspecting each variable’s frequency histogram and testing several values (Figure S9, Additional file 1). We found a suitable threshold for the number of missing observations to be around 25-30% of the number of individuals in the family, for mean site depth to be around 99th percentile, for mean genotype quality to be around 10th percentile, and for minor allele frequency to be either 0.38 or 0.10. Next, we phased the filtered genotypes using a haplotype doubling method [32] and converted the phased and filtered genotypes matrix to a format suitable for MSTmap [60]. For MSTmap, we used the distance_function kosambi and population_type DH for all the families and family-specific values for the parameters cutoff_p_value (between 10^−6^ and 10^−10^) and missing_threshold (either 0.25 or 0.30). The variant sites clustered into expected 16 linkage groups for six out of the seven families. However, one family had very few markers: only 389, while the other families had between 5,000 and 17,000 markers. We discarded the family with 389 markers and converted linkage groups from the remaining five families to ALLMAPS compatible format. Scripts used for linkage map creation and conversion to ALLMAPS format, including those from the steps below, are available online (see Availability of data and materials).

For Bionano optical maps, we first scaffolded the assembly using the hybrid scaffolding option of IrysView software (version 2.5.1) and using the aggressive preset. The process generated an XMAP file, among others, containing the contig connectivity information, which we converted to ALLMAPS compatible format using bionano2Allmaps.pl script [61]. We eliminated paths with less than four markers before running ALLMAPS.

We mapped RNA-seq reads to our assembly using default parameters of bwa-mem (version 0.7.17) [51] and eliminated reads that mapped to more than one location in the genome [62]. Next, we generated *ab initio* gene predictions using AUGUSTUS (version 3.2.3; --gff3=on -- species=fly) [63]. Next, we used AGOUTI (version 0.3.3-25-ga7e65d6) [64] to generate contig connectivity information from read mappings and *ab initio* gene predictions. Finally, we used a custom script to convert AGOUTI’s output to ALLMAPS compatible format.

## Supporting information

Table S1 - Assembly parameters

Table S2 - Contaminants identified in the best assembly

Table S3 - Comparison of the presented assembly with other fire ant genome assemblies

## Declaration

### Ethics approval and consent to participate

Not applicable.

## Consent for publication

Not applicable.

## Availability of data and materials

The Pacbio data that were used to generate the 36 assemblies as well as the scaffolded best assembly are available from NCBI (BioProject PRJNA609320).

The CompareGenomeQualities software is freely available under GPL-3.0 license from GitHub: https://github.com/wurmlab/CompareGenomeQualities. The software runs in the Unix command-line; we recommend using Bioconda (https://bioconda.github.io) or Docker (www.docker.com/) to install its dependencies (see Supplementary text, Additional file 1). The manuscript refers to commit c9aefc1 of the repository.

The set of scripts used to create linkage maps, and to convert linkage maps and contig connectivity information from Bionano and RNA-seq data to ALLMAPS compatible format are freely available under GPL-3.0 license: https://github.com/wurmlab/to_allmaps. The scripts are written in Bash, R, and Ruby programming languages. The manuscript refers to commit aef582d of the repository.

## Authors’ contributions

AP and YW conceived the experiment. ES sampled and genotyped the ants and extracted the DNA. AP performed the analysis, except the following: AW performed statistical tests for significance and AB conducted tests for foreign DNA contaminants. AP and YW wrote the manuscript. ES provided helpful comments on an initial draft of the manuscript. All authors subsequently contributed to improving the manuscript.

## Acknowledgements

We thank Maria Cristina Arias for providing the samples as permit holder; Andrew Leitch, Richard Nichols, Stephen Rossiter, James Borrell, and Marian Priebe for valuable discussions that has shaped the work; Simon Butcher, Chris Walker, Peter Childs, Dan Whitehouse, and Tom Bradford for their help with Queen Mary University of London’s Apocrita cluster (https://doi.org/10.5281/zenodo.438045); Rodrigo Pracana, Emeline Favreau and Ilya Levantis for comments on the manuscript; Philip Howard and Martin Tran for Bionano hybrid scaffolding; the UK’s JASMIN data analysis facility (https://www.jasmin.ac.uk).

## Additional files

Additional file 1 (DOCX): Supplementary text, Figures S1-S9, and Tables S4 and S5

Additional file 2 (.XLSX): Table S1 – Assembly parameters

Additional file 3 (.XLSX): Table S2 – Contaminants identified in the best assembly

Additional file 4 (.XLSX): Table S3 – Comparison of the presented assembly with other fire ant genome assemblies

## References

1. M Real F, Haas SA, Franchini P, Xiong P, Simakov O, Kuhl H, et al. The mole genome reveals regulatory rearrangements associated with adaptive intersexuality. Science. 2020;370:208–14.

2. Raymond O, Gouzy J, Just J, Badouin H, Verdenaud M, Lemainque A, et al. The Rosa genome provides new insights into the domestication of modern roses. Nat Genet. 2018;50:772–7.

3. Kronenberg ZN, Fiddes IT, Gordon D, Murali S, Cantsilieris S, Meyerson OS, et al. High-resolution comparative analysis of great ape genomes. Science. 2018;360:eaar6343.

4. Zhou Y, Shearwin-Whyatt L, Li J, Song Z, Hayakawa T, Stevens D, et al. Platypus and echidna genomes reveal mammalian biology and evolution. Nature. 2021;592:756–62.

5. Schatz MC, Delcher AL, Salzberg SL. Assembly of large genomes using second-generation sequencing. Genome Res. 2010;20:1165–73.

6. Ye L, Hillier LW, Minx P, Thane N, Locke DP, Martin JC, et al. A vertebrate case study of the quality of assemblies derived from next-generation sequences. Genome Biol. 2011;12:R31.

7. Alkan C, Sajjadian S, Eichler EE. Limitations of next-generation genome sequence assembly. Nat Methods. 2011;8:61–5.

8. Phillippy AM, Schatz MC, Pop M. Genome assembly forensics: finding the elusive mis-assembly. Genome Biol. 2008;9:R55.

9. Liu D, Hunt M, Tsai IJ. Inferring synteny between genome assemblies: a systematic evaluation. BMC Bioinformatics. 2018;19:26.

10. Denton JF, Lugo-Martinez J, Tucker AE, Schrider DR, Warren WC, Hahn MW. Extensive error in the number of genes inferred from draft genome assemblies. PLoS Comput Biol. 2014;10:e1003998.

11. Florea L, Souvorov A, Kalbfleisch TS, Salzberg SL. Genome assembly has a major impact on gene content: A comparison of annotation in two Bos taurus assemblies. PLoS One. 2011;6:e21400.

12. Vollger MR, Dishuck PC, Sorensen M, Welch AE, Dang V, Dougherty ML, et al. Long-read sequence and assembly of segmental duplications. Nat Methods. 2019;16:88–94.

13. Miga KH, Koren S, Rhie A, Vollger MR, Gershman A, Bzikadze A, et al. Telomere-to-telomere assembly of a complete human X chromosome. Nature. 2020;585:79–84.

14. Koren S, Harhay GP, Smith TPL, Bono JL, Harhay DM, Mcvey SD, et al. Reducing assembly complexity of microbial genomes with single-molecule sequencing. Genome Biol. 2013;14:R101.

15. Rhoads A, Au KF. PacBio sequencing and its applications. Genomics Proteomics Bioinformatics. 2015;13:278–89.

16. Mikheenko A, Prjibelski A, Saveliev V, Antipov D, Gurevich A. Versatile genome assembly evaluation with QUAST-LG. Bioinformatics. 2018;34:i142–50.

17. Kolmogorov M, Yuan J, Lin Y, Pevzner PA. Assembly of long, error-prone reads using repeat graphs. Nat Biotechnol. 2019;37:540–6.

18. Conte MA, Gammerdinger WJ, Bartie KL, Penman DJ, Kocher TD. A high quality assembly of the Nile Tilapia (Oreochromis niloticus) genome reveals the structure of two sex determination regions. BMC Genomics. 2017;18:341.

19. Minio A, Massonnet M, Figueroa-Balderas R, Castro A, Cantu D. Diploid genome assembly of the wine grape Carménère. G3: Genes, Genomes, Genetics. 2019;9:g3.400030.2019.

20. Zhang H, Jain C, Aluru S. A comprehensive evaluation of long read error correction methods. BMC Genomics. 2020;21:889.

21. Myers EW. A Whole-Genome Assembly of Drosophila. Science. 2000;287:2196–204.

22. Simão FA, Waterhouse RM, Ioannidis P, Kriventseva EV, Zdobnov EM. BUSCO: assessing genome assembly and annotation completeness with single-copy orthologs. Bioinformatics. Oxford University Press; 2015;31:3210–2.

23. Riba-Grognuz O, Keller L, Falquet L, Xenarios I, Wurm Y. Visualization and quality assessment of de novo genome assemblies. Bioinformatics. 2011;27:3425–6.

24. Khelik K, Sandve GK, Nederbragt AJ, Rognes T. NucBreak: Location of structural errors in a genome assembly by using paired-end Illumina reads. BMC Bioinformatics. 2020;21:393488.

25. Thomas GWC, Hahn MW. Referee: Reference Assembly Quality Scores. Genome Biol Evol. 2019;11:1483–6.

26. Koren S, Walenz BP, Berlin K, Miller JR, Bergman NH, Phillippy AM. Canu: scalable and accurate long-read assembly via adaptive -mer weighting and repeat separation. Genome Res. 2017;27:722–36.

27. Tschinkel WR. The Fire Ants. Harvard University Press; 2006.

28. Wurm Y, Wang J, Riba-Grognuz O, Corona M, Nygaard S, Hunt BG, et al. The genome of the fire ant Solenopsis invicta. Proc Natl Acad Sci U S A. 2011;108:5679–84.

29. Stolle E, Pracana R, Howard P, Paris CI, Brown SJ, Castillo-Carrillo C, et al. Degenerative expansion of a young supergene. Mol Biol Evol. 2019;36:553–61.

30. Pracana R, Levantis I, Martínez-Ruiz C, Stolle E, Priyam A, Wurm Y. Fire ant social chromosomes: differences in number, sequence and expression of odorant binding proteins. Evol Lett. 2017;1:199–210.

31. Venthur H, Zhou J-J. Odorant Receptors and Odorant-Binding Proteins as Insect Pest Control Targets: A Comparative Analysis. Front Physiol. 2018;9:1163.

32. Wang J, Wurm Y, Nipitwattanaphon M, Riba-Grognuz O, Huang Y-C, Shoemaker D, et al. A Y-like social chromosome causes alternative colony organization in fire ants. Nature. 2013;493:664–8.

33. Privman E, Wurm Y, Keller L. Duplication and concerted evolution in a master sex determiner under balancing selection. Proc Biol Sci. 2013;280:20122968.

34. Buechel SD, Wurm Y, Keller L. Social chromosome variants differentially affect queen determination and the survival of workers in the fire ant Solenopsis invicta. Mol Ecol. 2014;23:5117–27.

35. Pracana R, Priyam A, Levantis I, Nichols RA, Wurm Y. The fire ant social chromosome supergene variant Sb shows low diversity but high divergence from SB. Mol Ecol. 2017;26:2864–79.

36. Martinez-Ruiz C, Pracana R, Stolle E, Paris CI, Nichols RA, Wurm Y. Genomic architecture and evolutionary antagonism drive allelic expression bias in the social supergene of red fire ants. Elife. 2020;9:e55862.

37. Chin C-S, Alexander DH, Marks P, Klammer AA, Drake J, Heiner C, et al. Nonhybrid, finished microbial genome assemblies from long-read SMRT sequencing data. Nat Methods. 2013;10:563–9.

38. Roach MJ, Schmidt SA, Borneman AR. Purge Haplotigs: allelic contig reassignment for third-gen diploid genome assemblies. BMC Bioinformatics. 2018;19:460.

39. Liu B, Conroy JM, Morrison CD, Odunsi AO, Qin M, Wei L, et al. Structural variation discovery in the cancer genome using next generation sequencing: Computational solutions and perspectives. Oncotarget. 2015;6:5477–89.

40. Rahman A, Pachter L. CGAL: computing genome assembly likelihoods. Genome Biol. 2013;14:R8.

41. Logsdon GA, Vollger MR, Eichler EE. Long-read human genome sequencing and its applications. Nat Rev Genet. 2020;21:597–614.

42. Ballouz S, Dobin A, Gillis JA. Is it time to change the reference genome? Genome Biol. BioMed Central; 2019;20:1–9.

43. Walker BJ, Abeel T, Shea T, Priest M, Abouelliel A, Sakthikumar S, et al. Pilon: an integrated tool for comprehensive microbial variant detection and genome assembly improvement. PLoS One. 2014;9:e112963.

44. Rhie A, Walenz BP, Koren S, Phillippy AM. Merqury: reference-free quality, completeness, and phasing assessment for genome assemblies. Genome Biol. 2020;21:245.

45. Hunt GJ, Page RE Jr. Patterns of inheritance with RAPD molecular markers reveal novel types of polymorphism in the honey bee. Theor Appl Genet. 1992;85:15–20.

46. Li H. Minimap2: pairwise alignment for nucleotide sequences. Bioinformatics. 2018;34:3094–100.

47. Li H, Handsaker B, Wysoker A, Fennell T, Ruan J, Homer N, et al. The Sequence Alignment/Map format and SAMtools. Bioinformatics. 2009;25:2078–9.

48. Canu Parameter Reference. https://canu.readthedocs.io/en/latest/parameter-reference.html. Accessed 21 October 2017.

49. Berlin K, Koren S, Chin C-S, Drake JP, Landolin JM, Phillippy AM. Assembling large genomes with single-molecule sequencing and locality-sensitive hashing. Nat Biotechnol. 2015;33:623–30.

50. Gurevich A, Saveliev V, Vyahhi N, Tesler G. QUAST: quality assessment tool for genome assemblies. Bioinformatics. 2013;29:1072–5.

51. Li H. Aligning sequence reads, clone sequences and assembly contigs with BWA-MEM. 2013. 1303.3997.

52. Pedersen BS, Quinlan AR. Mosdepth: quick coverage calculation for genomes and exomes. Bioinformatics. 2018;34:867–8.

53. Quinlan AR, Hall IM. BEDTools: a flexible suite of utilities for comparing genomic features. Bioinformatics. 2010;26:841–2.

54. Pilon version 1.23. https://github.com/broadinstitute/pilon/releases/tag/v1.23. Accessed 24 August 2020.

55. Wood DE, Lu J, Langmead B. Improved metagenomic analysis with Kraken 2. Genome Biol. 2019;20:257.

56. Käfer S, Paraskevopoulou S, Zirkel F, Wieseke N, Donath A, Petersen M, et al. Re-assessing the diversity of negative strand RNA viruses in insects. PLoS Pathog. 2019;15:e1008224.

57. Calkins TL, Chen M-E, Arora AK, Hawkings C, Tamborindeguy C, Pietrantonio PV. Brain gene expression analyses in virgin and mated queens of fire ants reveal mating-independent and socially regulated changes. Ecol evol. 2018;8:4312–27.

58. Tang H, Zhang X, Miao C, Zhang J, Ming R, Schnable JC, et al. ALLMAPS: robust scaffold ordering based on multiple maps. Genome Biol. 2015;16:3.

59. Rochette NC, Rivera-Colón AG, Catchen JM. Stacks 2: Analytical methods for paired-end sequencing improve RADseq-based population genomics. Mol Ecol. 2019;28:4737–54.

60. Wu Y, Bhat PR, Close TJ, Lonardi S. Efficient and Accurate Construction of Genetic Linkage Maps from the Minimum Spanning Tree of a Graph. PLoS Genet. 2008;4:e1000212.

61. Zhang T. BioNano data revisited. https://github.com/tanghaibao/jcvi/issues/37#issuecomment-259032584. Accessed 6 June 2019.

62. Obtaining uniquely mapped reads from BWA mem alignment. https://bioinformatics.stackexchange.com/a/519. Accessed 12 June 2019.

63. Stanke M, Diekhans M, Baertsch R, Haussler D. Using native and syntenically mapped cDNA alignments to improve de novo gene finding. Bioinformatics. 2008;24:637–44.

64. Zhang SV, Zhuo L, Hahn MW. AGOUTI: improving genome assembly and annotation using transcriptome data. Gigascience. 2016;5:31.

