## Supplementary text, Figures S1-S9, and Tables S4 and S5 for "Parameter exploration improves the accuracy of long-read genome assembly"

Additional file 1

### Contents

#### Supplementary text

**CompareMyGenomes tool usage** **2**

**Comparison of Canu, flye, and wtdbg2 genome assemblers** **3**

**Quality control of Illumina reads** **4**

#### Supplementary figures

**Figure S1**: Length distribution of PacBio reads **5**

**Figure S2**: Coverage vs length plots for the best assembly **6**

**Figure S3**: Correlation between the four metrics: NG50, BUSCO score, resolved length, and solid Illumina read pairs **7**

**Figure S4**: Dotplot of the presented and the draft fire ant genome assembly **8**

**Figure S5**: Estimated error rate of corrected reads **9**

**Figure S6**: Coverage density plot of the best assembly before and after removing unresolved haplotypes **10**

**Figure S7**: R general linear model output – determining significance of assembly parameters **11**

**Figure S8**: Proportion of genotyped vs homozygous individuals from RAD-seq of seven fire ant families **12**

**Figure S9**: Histogram of the number of genotyped individuals, mean read depth, mean genotype quality and minor allele frequency obtained from RAD-seq of seven ant families. **13**

#### Supplementary tables

**Table S1:** Additional file 2

**Table S2:** Additional file 3

**Table S3:** Additional file 4

**Table S4**: Polishing and haplotype removal improves assembly accuracy **15**

**Table S5**: Comparison of Canu, Flye, and wtdbg2 genome assemblers **15**

#### References

### Supplementary text

**CompareGenomeQualities tool usage**

Our tool makes use of multiple programming languages and bioinformatics software [1]. To facilitate usage, we provide a bash script that can install the required dependencies using Bioconda [2]. We also provide our tool and all its dependencies as a Docker image [3]. Below, we first provide an overview of our tool's command-line parameters. We then present example usage of the tool.

<u>Overview of command-line parameters</u>

```
-b, --busco-lineage     One of the 193 BUSCO v5 datasets listed here:
                        https://busco-data.ezlab.org/v5/data/lineages. The
                        dataset is automatically downloaded. Dataset name
                        can be a partial match e.g., insecta instead of
                        insecta_odb10.2020-09-10.tar.gz. Required unless
                        --rank-only is specified.
-g, --genome-size       Expected or estimated genome size in base pairs.
                        Required unless --rank-only is specified.
-1, --illumina-R1       Forward Illumina reads. Required unless --rank-only
                        is specified.
-2, --illumina-R2       Reverse Illumina reads. Required unless --rank-only
                        is specified.
-n, --num-cpus          Used for read mapping and BUSCO steps. Default: 1.
-o, --output-dir        Output directory. Not applicable if --rank-only is
                        specified. Default: /mnt/compare-genome-qualities-
                        yyyy-mm-dd-hhmmss.
--rank-only             Don't compute metrics. Only rank assemblies based
                        on tabular files in the given directory.
-h, --help               View this message docker run
```

Example usage of the tool

There are two ways to run the tool. The default behavior is to run the tool on a series of genome assemblies, providing a set of Illumina reads as additional input. This will compute the assembly quality metrics NG50, BUSCO score, resolved length, and solid Illumina read pairs and subsequently rank the assemblies.

```
compare-genome-qualities.sh -g 450000000 -b insecta_odb9 -1
illumina_R1.fq.gz -2 illumina_R2.fq.gz assembly_1.fa assembly_2.fa
assembly_3.fa
```

Alternatively, our tool can be used to rank genome assemblies based on pre-computed metrics. The pre-computed metrics are presented to the tool in form of tabular files, one file per metric, each file containing one line per assembly indicating the assembly identifier and the value of the metric for that assembly, separated by the tab character.

```
compare-genomes-qualities.sh --rank-only dir_containing_tabular_files
```

**Comparison of Canu, flye, and wtdbg2 genome assemblers**

Our tool does not require the genome assemblies to be generated using different parameter combinations. For example, we present a comparison of the assemblies generated by three different long-read genome assembly tools: Canu [4], flye [5], and wtdbg2 [6] (Table S5). All assemblies were generated using default parameters of the assembly software. We removed unresolved haplotigs from Canu assembly [7] to get a better sense of resolved assembly lengths, but we did not polish any of the assemblies. In this example, wtdbg2 generated the most contiguous assembly. However, the assembly generated by Canu had the most resolved regions (13 Mb more than the next best) and considerably higher proportion of solidly mapped Illumina reads (57.62% compared to 55.25% of the runner up), followed by Flye. The 0.01% difference in the BUSCO score [8] of Canu and Flye assemblies is minor and likely to be eliminated by subsequent polishing steps. These results validate our choice of using Canu for assembly parameter optimization and further highlight the benefits of testing different assembly software for a given dataset.

### Quality control of Illumina reads

We filtered and trimmed Illumina datasets prior to use. First, we removed optical duplicates using clumpify.sh (version 37) [9]. Second, we removed reads with mean quality threshold lower than 15 using htqc [10]. Third, we compared the mean base quality per cycle, per tile to the mean base quality of that cycle across all tiles to test for air-bubbles becoming trapped in the flow cell [11]. For this, we obtained the difference between per-cycle mean base quality for a tile and the per-cycle mean base quality for all tiles from FastQC's text output (version 0.11.5) [12]. Where this difference was greater than 4, we changed the corresponding base in the reads to 'N'. This was done by creating a BED file of positions from the tile and cycle information and then using seqtk (version 1.2) [13] to convert bases at the positions specified in the file. Next, we considered that reads with multiple occurrences of low-quality bases may be problematic. To eliminate such reads, we turned bases with quality scores lower than 12 to 'N' using seqtk (reads with excessive Ns are removed in the next step). Finally, we used cutadapt (version 1.13) [14] to trim adapter sequences, to trim low quality bases from both the 3' and 5' ends, to trim any leading and trailing 'N's, to eliminate after trimming reads shorter than 50 bp and those with more than 4 'N's. For the Illumina sequences used for assembly comparison, we retained 64,850,542 pairs of 50-150 bp reads (*i.e.*, 79.23% of sequenced bases) after filtering.

### Supplementary figures

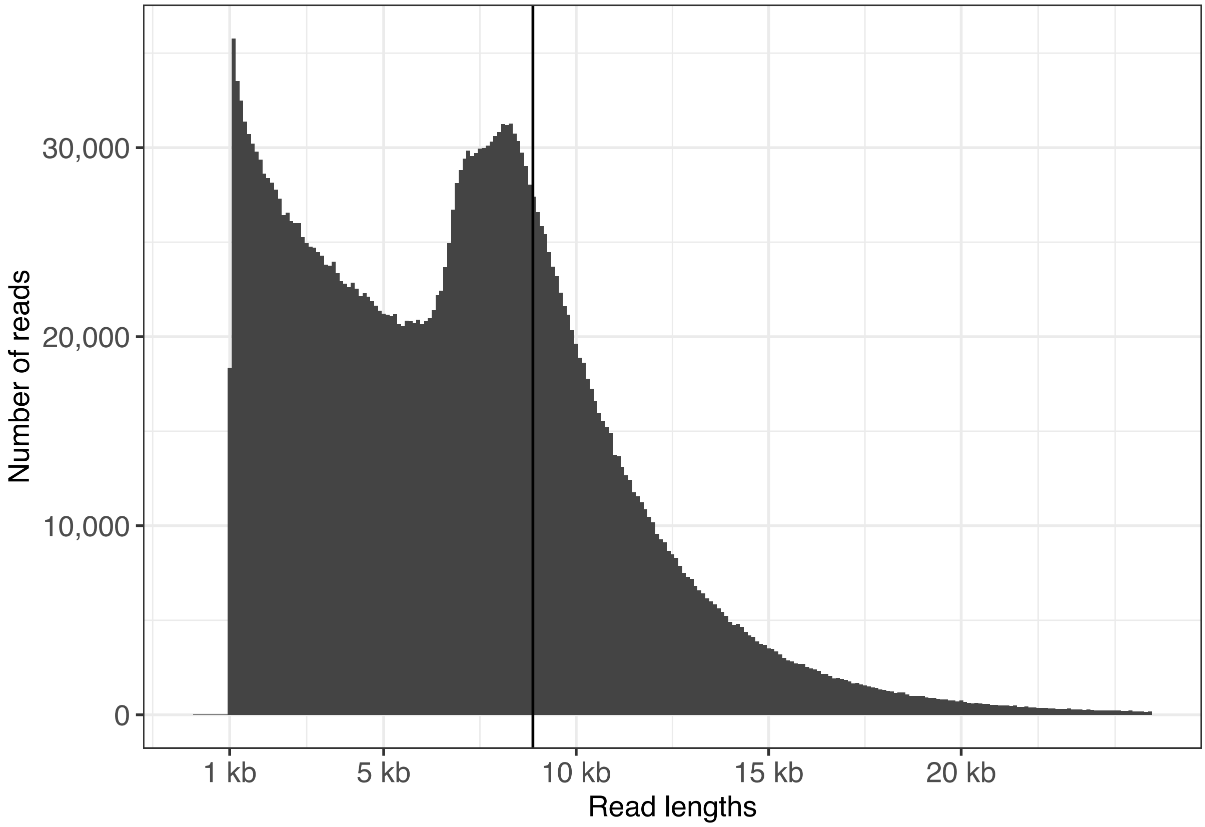


**Figure S1**: Histogram of lengths of raw Pacbio reads. The black vertical line shows N50 read length (8,876 bp).

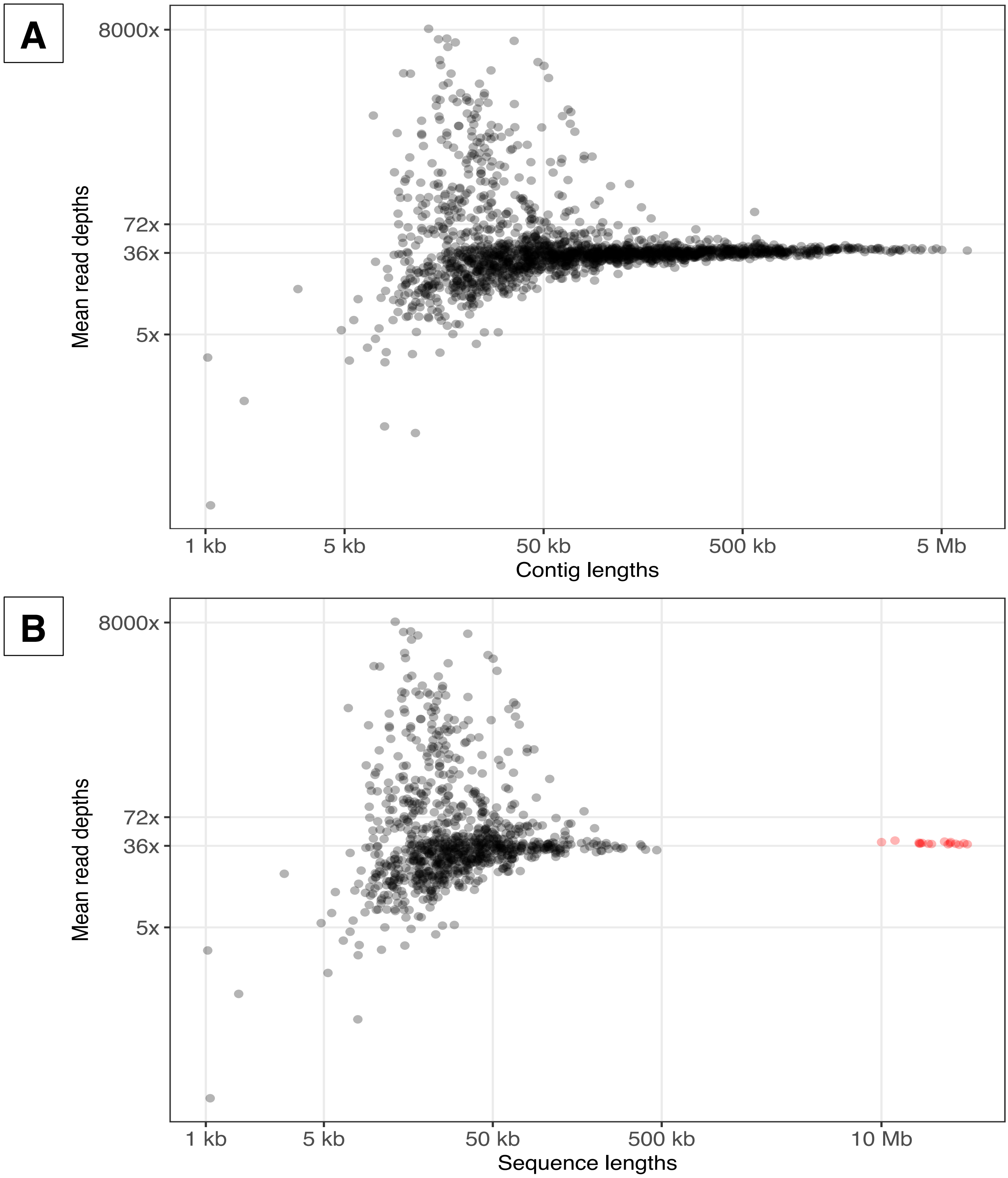


**Figure S2**: Lengths of the assembled sequences of the best assembly (x axis) and their average read depths (y axis) A) before scaffolding B) and after. In panel B, the sequences longer than 10 Mb (colored red) are the chromosomes, while the cloud of sequences on the left are unplaced contigs. Axes are log scaled. Pacbio reads were mapped to the best assembly using minimap2 (version 2.17; -a -x map-pb) [15]. Read depth of contigs were calculated using mosdepth (version 0.2.6; -x -n) [16]. Contigs with average depth higher than twice the median coverage (36x) are likely to contain collapsed representation of highly repetitive regions of the genome. Contigs with average depth lower than 5x are likely to contain higher amounts of sequencing error. This is because the SMRTLinks polishing step, which is critical for Canu assemblies, excludes regions with coverage lower than 5x.

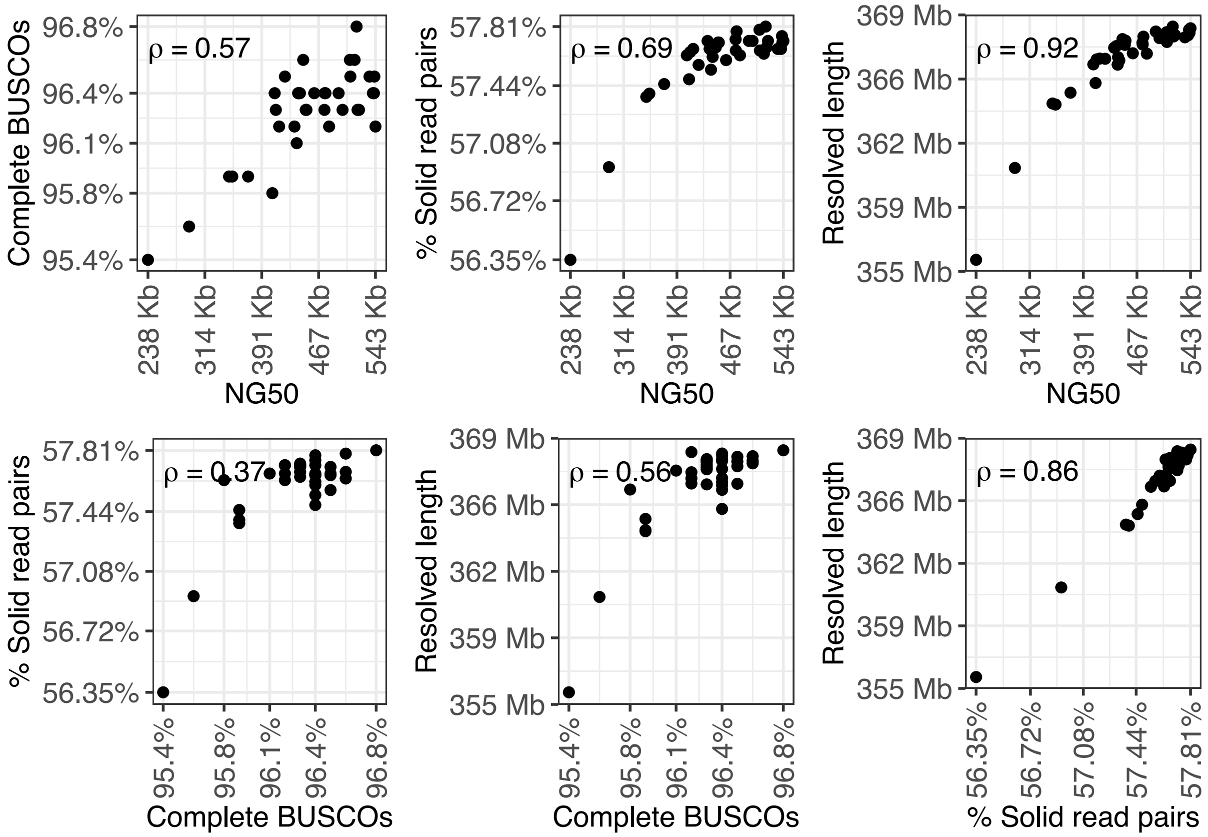


**Figure S3**: Correlations between the four metrics of genome quality: NG50, BUSCO score, resolved length, and solid Illumina read pairs. Each panel shows the values taken by a pair of metrics on the x and the y axes, and Spearman's rank correlation coefficient (ρ) between the metrics. To account for the general correlation across metrics, the overall ranking of assemblies performed by CompareGenomeQualities is weighted by the complement of the average pairwise correlations (Fig 1).

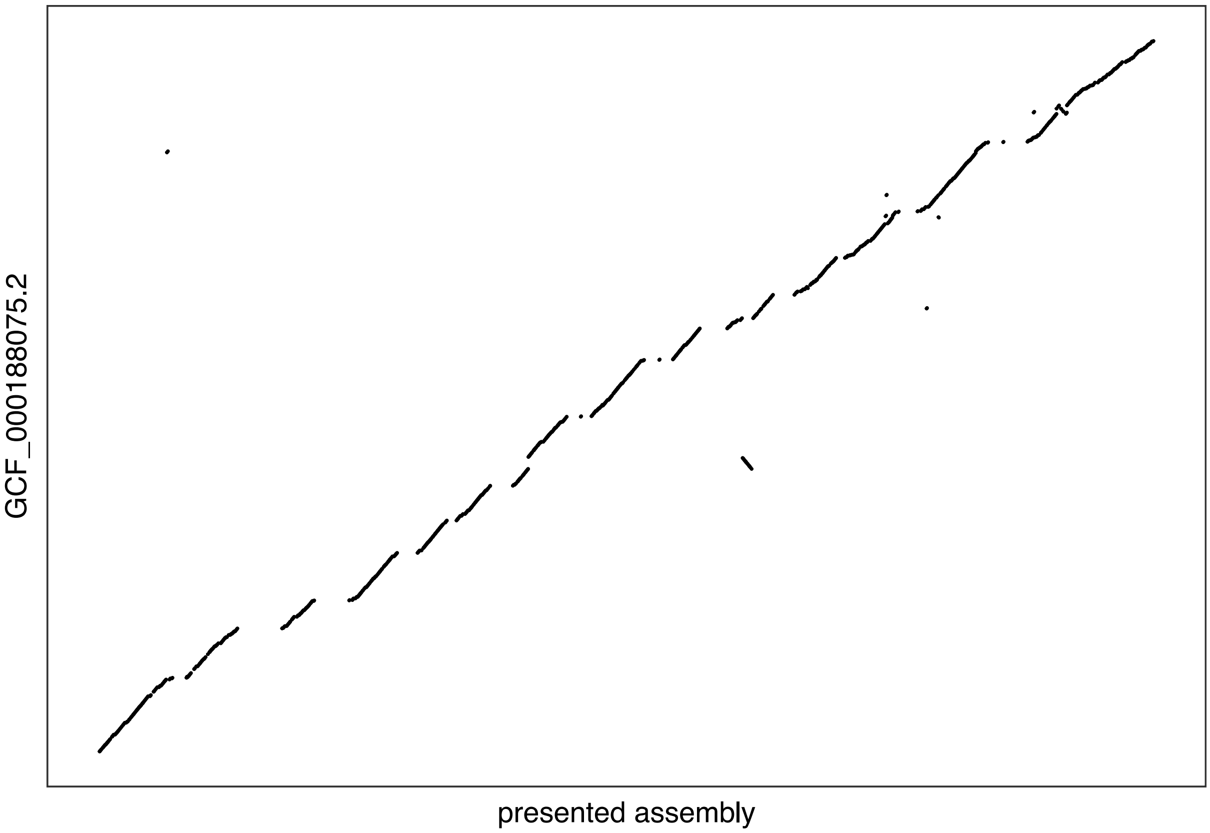


**Figure S4**: Dotplot comparison of the fire ant genome assembly we present here (x axis) against a previously published draft fire ant genome [17] (y axis). The x axis represents the 16 fire ant chromosomes in the presented assembly and the y axis represents matching sequences in the draft assembly. The assemblies were aligned using minimap2 (version 2.17; -c -P -k19 -w19 -m200) [15] and visualized using dotPlotly (version 11744849; -m 100000) [18]. Most breaks in collinearity are along the x axis. The spacing between diagonals shows how ambiguous regions of the genome that are absent from the previous genome assembly were identified and included in the new assembly.

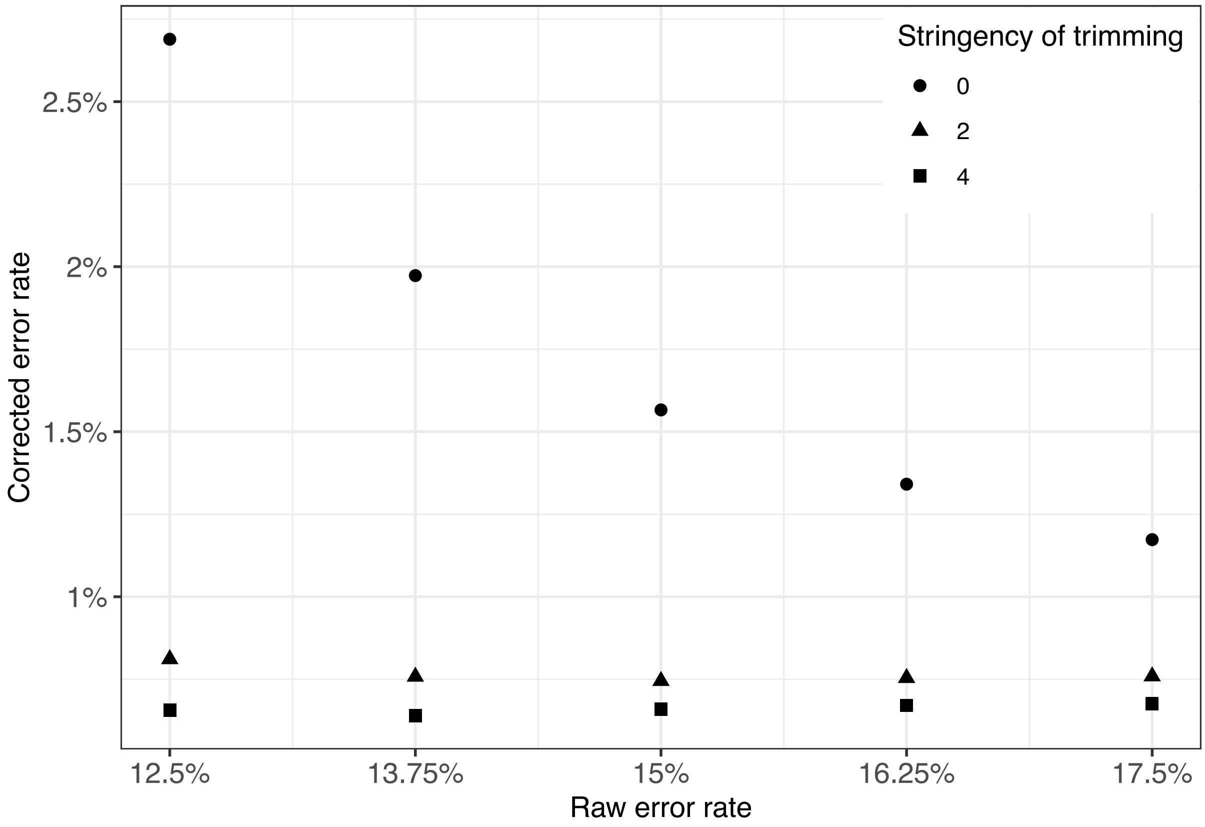


**Figure S5**: Estimated error rates of corrected reads (y axis) against error rate of raw reads (x axis). Shape of the points indicate the stringency of trimming raw reads. Each of the 15 points represents corrected reads obtained by changing the raw error rate threshold and trimming stringency genome assembly parameters used for this study.

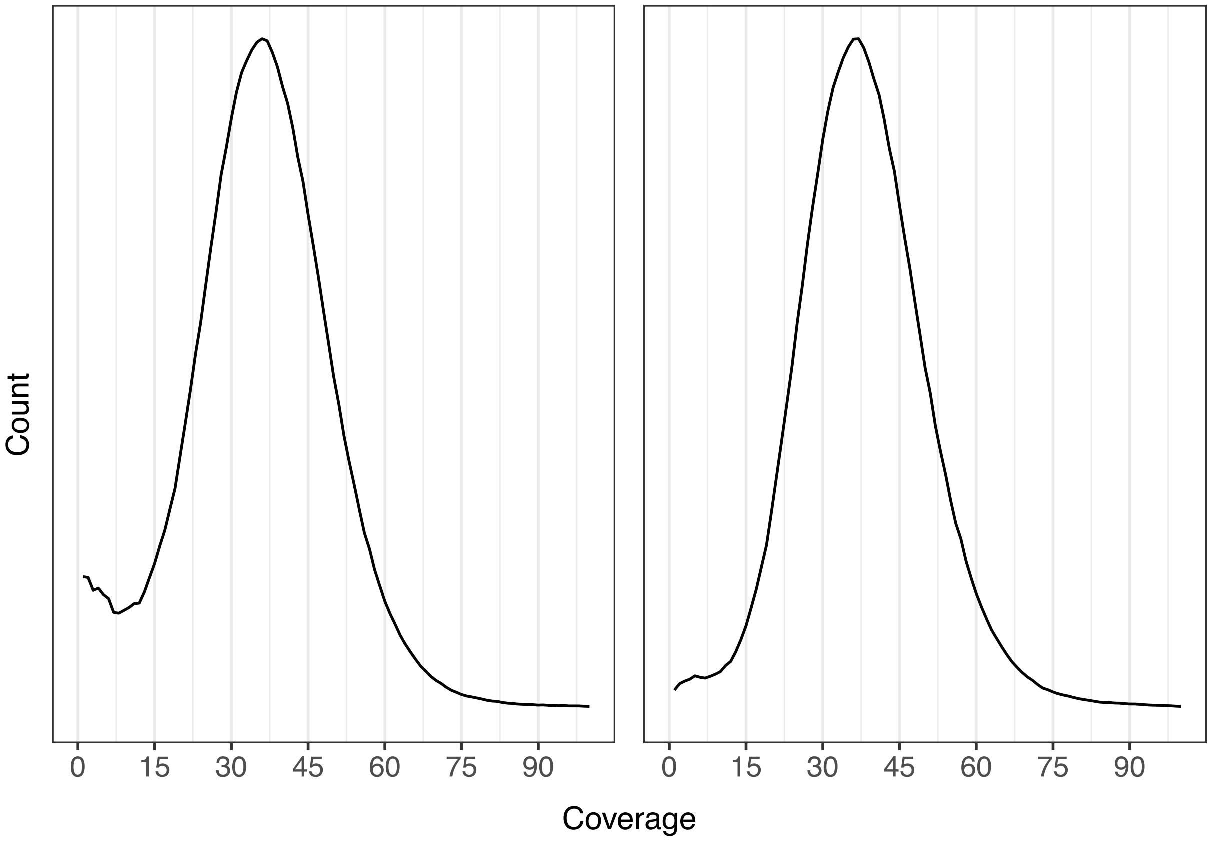


**Figure S6**: Coverage density plot of the best assembly before (left) and after removing unresolved haplotigs (right). The removal of these unresolved haplotigs clearly reduces the among of contigs with less than half the median coverage (*i.e.*, less than 18x).

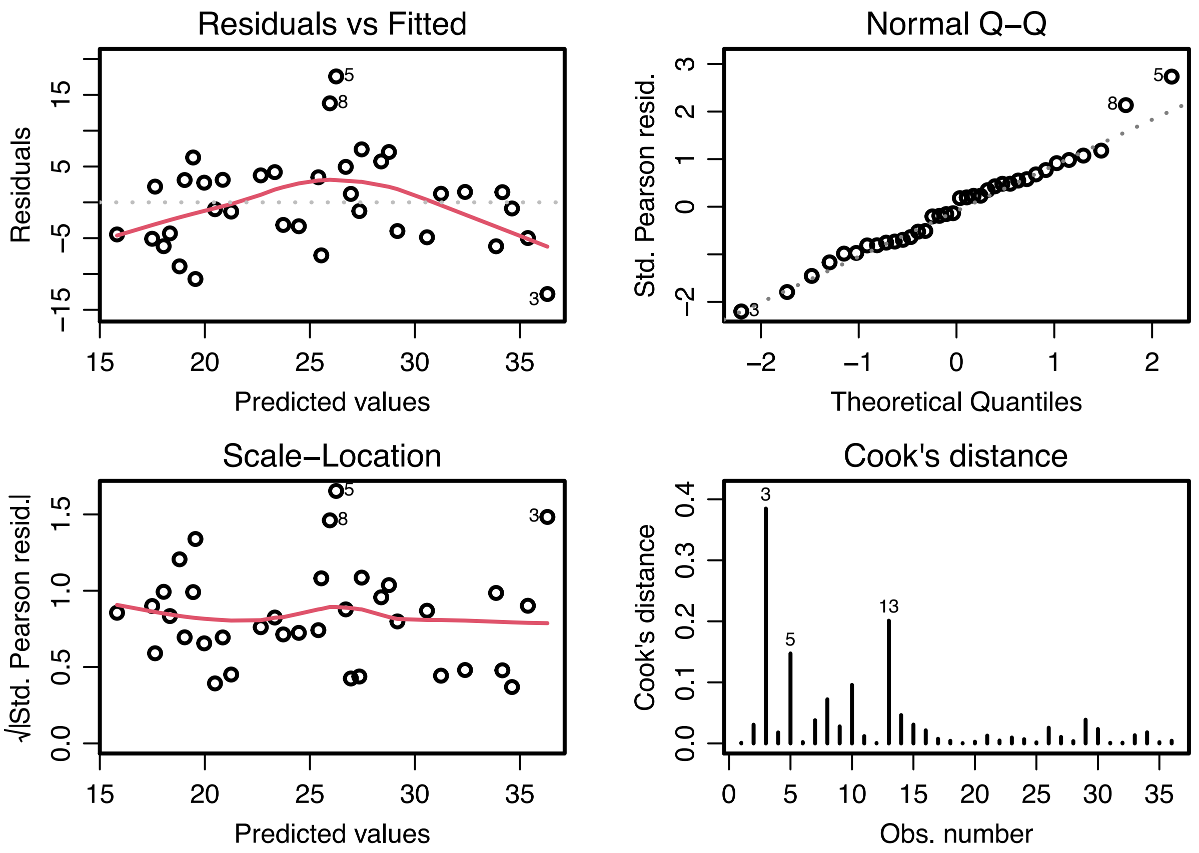


**Figure S7**: We modelled the overall assembly rank as a function of the three assembly parameters: error rate threshold for raw reads, stringency of trimming raw reads, error rate threshold for corrected reads). The error rate threshold for corrected and for raw reads were significant $p < 10^{-5}$ and $p < 0.05$ respectively.

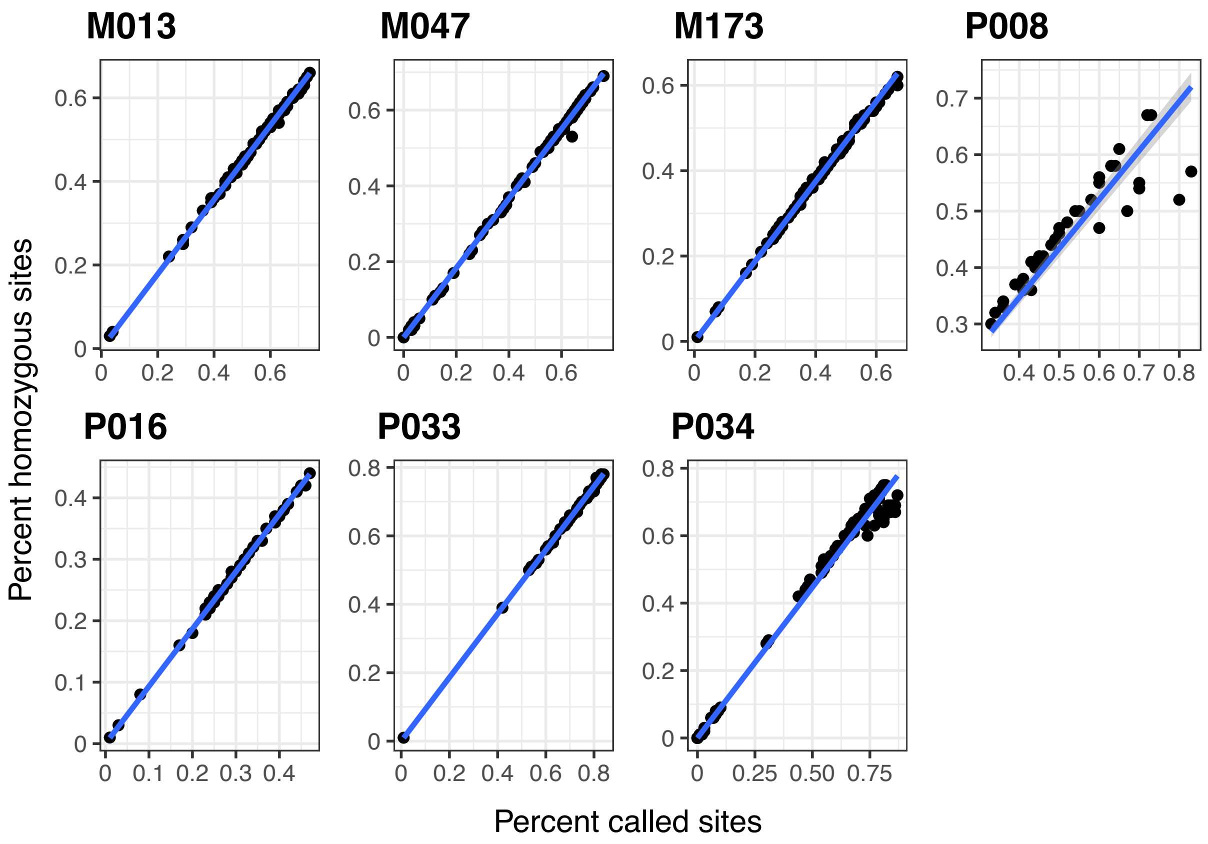


**Figure S8**: Proportion of individuals genotyped per site from RAD sequencing of seven fire ant families (M013 - P034; names beginning with M are monogynous colonies and those beginning with P are polygynous colonies) that has single nucleotide polymorphism in the family (x axis) against proportion of homozygous individuals for that site (y axis).

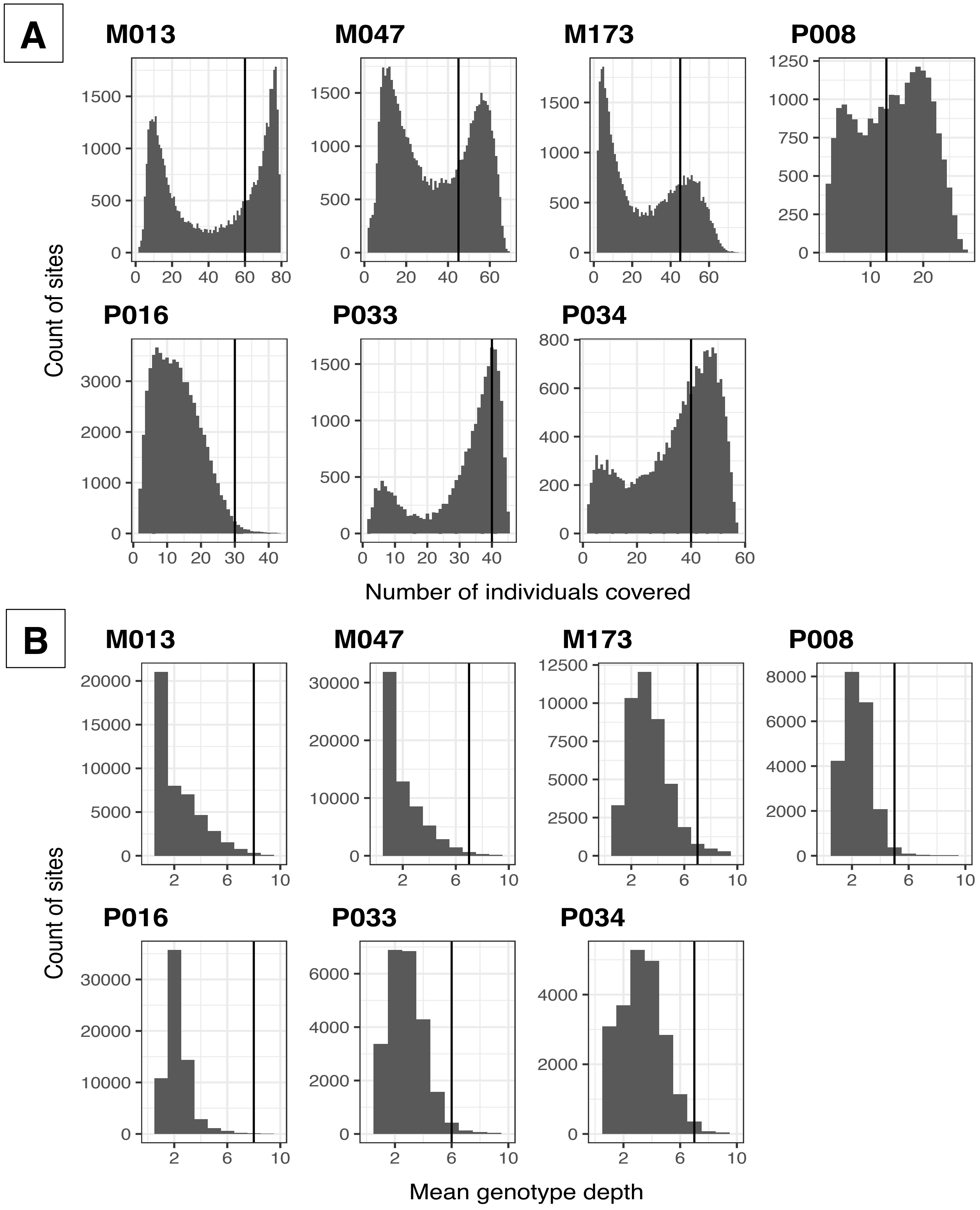


**Figure S9**: Site statistics obtained from RAD sequencing of seven fire ant families (M013 - P034; names beginning with M are monogynous colonies and those beginning with P are polygynous colonies). Black vertical line shows the threshold chosen for each family for filtering during linkage map construction. A) Number of individuals genotyped per site (x axis)

against their count (y axis). B) Mean read depth of genotypes per site (x axis) against their count (y axis). C) Mean genotype quality per site (x axis) against their count (y axis). D) Minor allele frequency per site (x axis) against their count (y axis) (continued on next page)

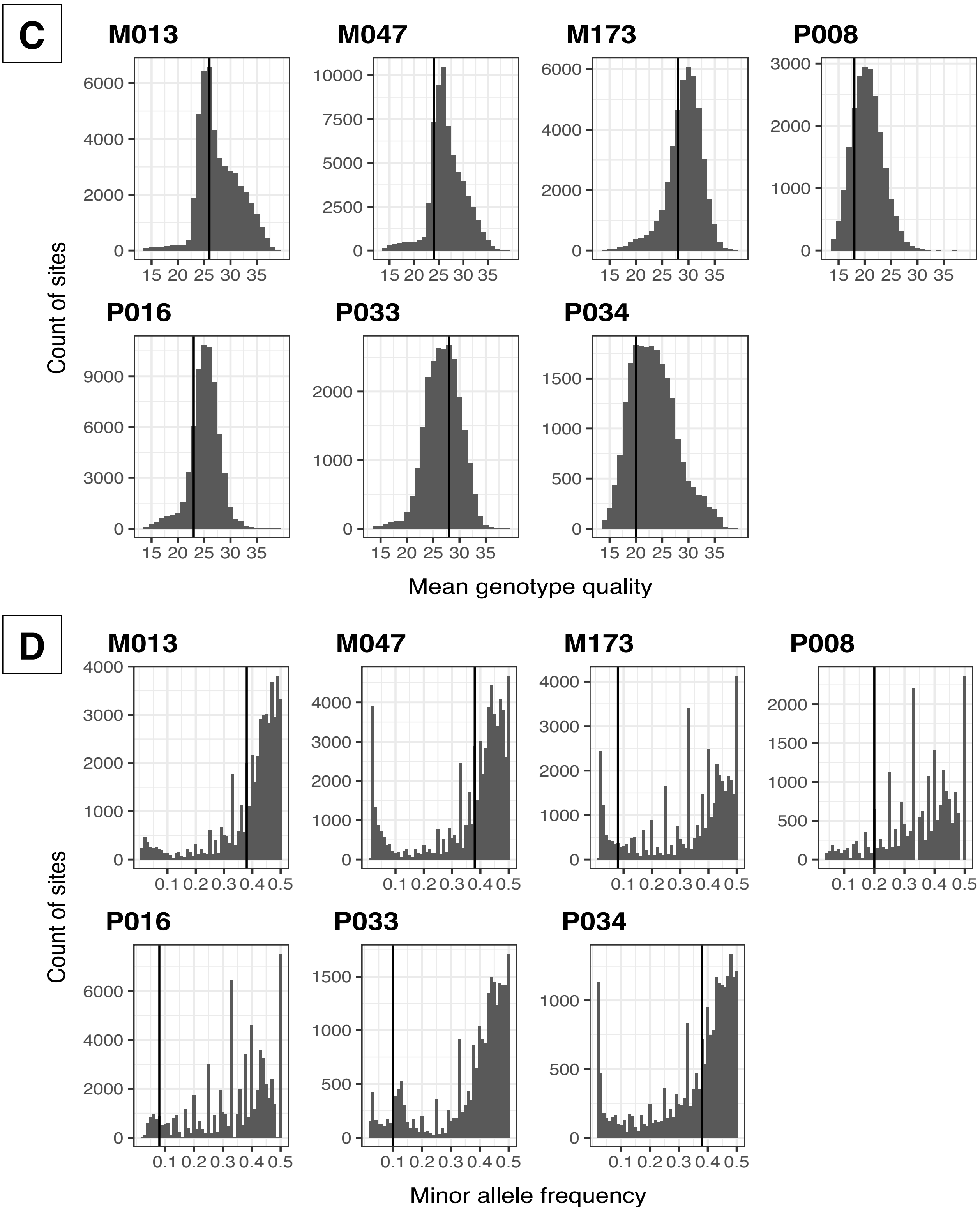


**Figure S9 (continued)**

### Supplementary tables

**Table S1:** Additional file 2

**Table S2:** Additional file 3

**Table S3:** Additional file 4

**Table S4**: Polishing and haplotype removal improves assembly accuracy

| | Raw contigs | After Pacbio polishing | After further Illumina polishing | After further haplotype-filtering |
|---|---|---|---|---|
| General error rate | 1.34 | 1.30 | 1.26 | - |
| % Illumina reads with insertion | 6.48% | 3.91% | 1.79% | - |
| % Illumina reads with deletion | 2.75% | 2.82% | 2.19% | - |
| Mean mapping quality of Illumina reads | 25.73 | 27.49 | 28.24 | - |
| % Complete benchmarking universal single-copy orthologs (BUSCO, lineage insecta, n=1664) | 98% | 98.8% | 99% | 98.8% |

**Table S5**: Comparison of Canu, Flye, and wtdbg2 genome assemblers

| | Resolved length | NG50 | BUSCO score | Solid Illumina read pairs |
|---|---|---|---|---|
| Canu + purge_haplotigs | 366,814,754 bp | 441,945 bp | 96.4% | 57.62% |
| Flye | 353,678,069 bp | 402,671 bp | 96.5% | 55.25% |
| Wtdbg2 | 320,860,502 bp | 502,081 bp | 68.6% | 48.12% |

### References

1. CompareGenomeQualities. https://github.com/wurmlab/CompareGenomeQualities. Accessed 14 May 2021.

2. Grüning B, Dale R, Sjödin A, Chapman BA, Rowe J, Tomkins-Tinch CH, et al. Bioconda: sustainable and comprehensive software distribution for the life sciences. Nat Methods. 2018;15:475–6.

3. Empowering App Development for Developers. https://www.docker.com/. Accessed 11 May 2021.

4. Koren S, Walenz BP, Berlin K, Miller JR, Bergman NH, Phillippy AM. Canu: scalable and accurate long-read assembly via adaptive -mer weighting and repeat separation. Genome Res. 2017;27:722–36.

5. Kolmogorov M, Yuan J, Lin Y, Pevzner PA. Assembly of long, error-prone reads using repeat graphs. Nat Biotechnol. 2019;37:540–6.

6. Ruan J, Li H. Fast and accurate long-read assembly with wtdbg2. Nat Methods. 2020;17:155–8.

7. Roach MJ, Schmidt SA, Borneman AR. Purge Haplotigs: allelic contig reassignment for third-gen diploid genome assemblies. BMC Bioinformatics. 2018;19:460.

8. Simão FA, Waterhouse RM, Ioannidis P, Kriventseva EV, Zdobnov EM. BUSCO: assessing genome assembly and annotation completeness with single-copy orthologs. Bioinformatics. 2015;31:3210–2.

9. Bushnell B. Introducing Clumpify. https://www.biostars.org/p/225338/. Accessed 5 June 2017.

10. Yang X, Liu D, Liu F, Wu J, Zou J, Xiao X, et al. HTQC: a fast quality control toolkit for Illumina sequencing data. BMC Bioinformatics. 2013;14:33.

11. Andrews S. Position specific failures of flowcells. https://sequencing.qcfail.com/articles/position-specific-failures-of-flowcells/. Accessed 9 June 2017.

12. FastQC: A quality control tool for high throughput sequence data. https://www.bioinformatics.babraham.ac.uk/projects/fastqc. Accessed 11 Feb 2017.

13. Li H. seqtk: Toolkit for processing sequences in FASTA/Q formats. https://github.com/lh3/seqtk. Accessed 15 June 2017.

14. Martin M. Cutadapt removes adapter sequences from high-throughput sequencing reads. EMBnet.journal. 2011;17:10–2.

15. Li H. Minimap2: pairwise alignment for nucleotide sequences. Bioinformatics. 2018;34:3094–100.

16. Pedersen BS, Quinlan AR. Mosdepth: quick coverage calculation for genomes and exomes. Bioinformatics. 2018;34:867–8.

17. Wurm Y, Wang J, Riba-Grognuz O, Corona M, Nygaard S, Hunt BG, et al. The genome of the fire ant *Solenopsis invicta*. Proc Natl Acad Sci U S A. 2011;108:5679–84.

18. Poorten T. dotPlotly: Generate an interactive dot plot from mummer or minimap alignments. https://github.com/tpoorten/dotPlotly. Accessed 23 December 2020.
